# Somatic inactivating *PTPRJ* mutations and dysregulated pathways identified in canine melanoma by integrated comparative genomic analysis

**DOI:** 10.1101/196337

**Authors:** W Hendricks, V Zismann, K Sivaprakasam, C Legendre, K Poorman, W Tembe, J Kiefer, W Liang, V DeLuca, M Stark, A Ruhe, R Froman, N Duesbury, M Washington, Jessica Aldrich, M Neff, M Huentelman, N Hayward, K Brown, D Thamm, G Post, C Khanna, B Davis, M Breen, Aleksandar Sekulic, J Trent

## Abstract

Canine malignant melanoma, a significant cause of mortality in domestic dogs, is a powerful comparative model for human melanoma, but little is known about its genetic etiology. We mapped the genomic landscape of canine melanoma through multi-platform analysis of 37 tumors (31 mucosal, 3 acral, 2 cutaneous, and 1 uveal) and 17 matching constitutional samples including long- and short-insert whole genome sequencing, RNA sequencing, array comparative genomic hybridization, single nucleotide polymorphism array, and targeted Sanger sequencing analyses. We identified novel predominantly truncating mutations in the putative tumor suppressor gene *PTPRJ* in 19% of cases. No *BRAF* mutations were detected, but activating *RAS* mutations (24% of cases) occurred in conserved hotspots in all cutaneous and acral and 13% of mucosal subtypes. *MDM2* amplifications (24%) and *TP53* mutations (19%) were mutually exclusive. Additional low-frequency recurrent alterations were observed amidst low point mutation rates, an absence of ultraviolet light mutational signatures, and an abundance of copy number and structural alterations. Mutations that modulate cell proliferation and cell cycle control were common and highlight therapeutic axes such as MEK and MDM2 inhibition. This mutational landscape resembles that seen in *BRAF* wild-type and sun-shielded human melanoma subtypes. Overall, these data inform biological comparisons between canine and human melanoma while suggesting actionable targets in both species.

**AUTHOR SUMMARY:** Melanoma, an aggressive cancer arising from transformed melanocytes, commonly occurs in pet dogs. Unlike human melanoma, which most often occurs in sun-exposed cutaneous skin, canine melanoma typically arises in sun-shielded oral mucosa. Clinical features of canine melanoma resemble those of human melanoma, particularly the less common sun-shielded human subtypes. However, whereas the genomic basis of diverse human melanoma subtypes is well understood, canine melanoma genomics remain poorly defined. Similarly, although diverse new treatments for human melanoma based on a biologic disease understanding have recently shown dramatic improvements in outcomes for these patients, treatments for canine melanoma are limited and outcomes remain universally poor. Detailing the genomic basis of canine melanoma thus provides untapped potential for improving the lives of pet dogs while also helping to establish canine melanoma as a comparative model system for informing human melanoma biology and treatment. In order to better define the genomic landscape of canine melanoma, we performed multi-platform characterization of 37 tumors. Our integrated analysis confirms that these tumors commonly contain mutations in canine orthologs of human cancer genes such as *RAS*, *MDM2*, and *TP53* as well mutational patterns that share important similarities with human melanoma subtypes. We have also found a new putative cancer gene, *PTPR*J, frequently mutated in canine melanoma. These data will guide additional biologic and therapeutic studies in canine melanoma while framing the utility of comparative studies of canine and human cancers more broadly.

## INTRODUCTION

Human melanoma is of increasing clinical concern. It is one of a few cancers with rising incidence, while five-year survival for patients with metastatic disease has until recently remained low (15-20%) due to a dearth of curative systemic therapies(1). Discovery of frequent activating BRAF mutations in melanoma and treatment with selective inhibitors of this mutant kinase has led to dramatic responses in the setting of metastatic disease(2-4). However, not all *BRAF*-mutant melanomas respond to targeted therapy and responses that do occur are often brief and followed by the emergence of drug-resistant disease(5). Moreover, targeted treatment options in melanoma subtypes without activating *BRAF* mutations are limited. New treatment paradigms such as immunotherapy, drug combinations, and alternative dosing strategies may circumvent resistance and broaden the scope of precision medicine in melanoma(6-9), but rapid preclinical study of such regimens requires access to robust models that recapitulate complex tumor features such as intratumoral genomic heterogeneity and tumor-host interactions. Meanwhile, few animal models exist for uncommon molecular or histological melanoma subtypes such as *BRAF* wild-type (*BRAF*wt) or mucosal melanoma.

Naturally-occurring canine cancers are increasingly recognized as meeting a need for complex cancer models that develop gradually amidst interactions with host stroma and immune system(10-16). Spontaneous canine malignant melanomas, which are almost universally *BRAF*wt and for which the mucosal subtype is the most prevalent clinically significant form, may fill a specific gap in models of *BRAF*wt and rare histological melanoma subtypes(11). Human mucosal melanoma is an aggressive histological subtype that is predominantly *BRAF*, *RAS*, and *NF1* wild type (Triple Wild Type or TWT) with occasional mutations in *KIT* or *NRAS* and carries a five-year survival rate between 12.3% and 35.3% (17-26). Study of this subtype is limited by its low prevalence, only 1-2% of human melanomas in the United States, with as few as 1,500 cases per year(27). On the other hand, canine malignant melanoma accounts for up to 100,000 yearly cancer diagnoses in the United States, occurring most commonly in the oral mucosa, but also arising in cutaneous and acral epithelium(28-31).

Canine malignant melanoma is highly prevalent, closely mirrors human melanoma clinically and pathologically, and is extremely aggressive, with median survival for oral cases being a mere 200 days(32-36). However, little is known about its genetic etiology. It is predominantly *BRAF*wt with frequent copy number alterations of regions of canine chromosomes (CFA) 13, 17, 22, and 30, alongside frequent *MYC* amplifications and deletions of *CDKN2A*. Targeted sequencing studies, though limited, have shown that it infrequently bears alterations in other known drivers of human melanoma[32, 36-42]. It has been shown that CFA 30 aberrations are characteristic of canine oral melanoma and complex copy number profiles on this chromosome homologous to the same profiles on human chromosome (HSA) 15 in human mucosal melanoma are suggestive of rearrangements that may drive this melanoma subtype (41). Despite the very low prevalence of *BRAF* mutations, immunohistochemistry (IHC) has shown that the mitogen-activated protein kinase (MAPK) and/or phosphoinositide 3-kinase (PI3K) pathways are activated in 52-77% of cases[32, 36-40]. These data hint at underlying mutations driving these pathways that could guide future biological exploration and therapeutic development in the canine and human diseases.

We therefore set out to map the genomic landscape of canine melanoma using a combination of massively parallel whole genome sequencing (WGS), array-based platforms and targeted sequencing to identify somatic changes driving these cancers. Here we report the identification of recurrent inactivating mutations in the candidate tumor suppressor gene *PTPRJ* in addition to frequent *RAS* mutations, and mutually-exclusive *MDM2* and *TP53* alterations. We thereby define the genomic landscape of these cancers and identify similarities between melanoma subtypes across species while highlighting subtype-specific aberrations that may be used to guide future research.

## RESULTS

### Patterns of mutation identified by whole genome analysis of canine melanoma

We undertook comprehensive analysis of acquired genetic alterations in a discovery cohort of seven melanomas and matched germlines from six dogs (two tumors were derived from one dog) using WGS for detection of subtle sequence alterations alongside long-insert WGS (LI-WGS, see Materials and Methods)(43) for sensitive detection of structural variants. We then performed copy number and targeted gene analyses in an additional 27 tumors and three melanoma cell lines **(Table 1).** Tumors (all primary tumors except one acral metastasis) and matching whole blood were collected through the Van Andel Research Institute from dogs undergoing surgery at specialty veterinary clinics and immediately snap frozen. Diagnosis of melanoma was confirmed by two independent board certified veterinary pathologists in addition to staining for three melanocytic differentiation markers where tissue was available(36, 44). Diverse breeds are represented in this cohort with enrichment for Cocker Spaniels and Golden Retrievers (five dogs of each breed), an equal ratio of male and female dogs and a median age at resection of 11 years. Clinicopathologic characteristics for this cohort are described in **Supplementary Table 1** and **Supplementary Figure 1.**

**Table 1.** Summary of Genomic Analyses Performed in Canine Melanoma

| Analysis platform | Type of alteration detected | Samples analyzed |
| --- | --- | --- |
| Discovery cohort |  |  |
| WGS | Point mutations, copy number, structural alterations | 7 tumor and 6 matching normal samples |
| LI-WGS | Copy number and structural alterations |  |
| mRNASeq | Expressed point mutations and gene fusions |  |
| aCGH | Copy number alterations |  |
| SNP-A | Copy number alterations |  |
| Prevalence cohort |  |  |
| Targeted Sequencing | Point mutations | 27 tumor and 11 matching normal samples, 3 cell lines |
| SNP-A | Copy number alterations |  |
| Total distinct samples |  | 34 tumor samples, 18 matching normals, 3 cell lines |

**Figure 1.**
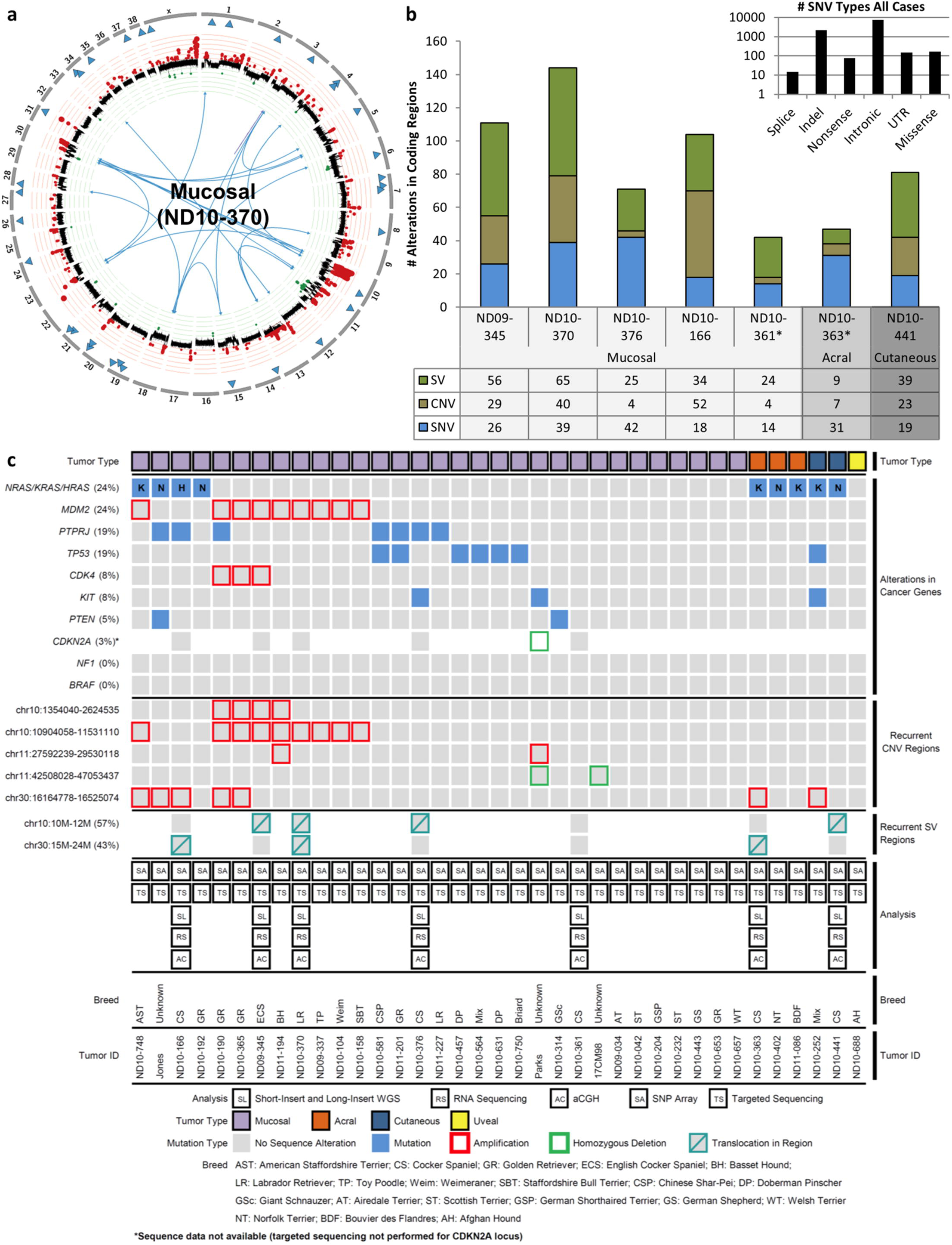
The mutational landscape of canine melanoma. (a) A representative Circos plot depicting coding SNVs, CNVs, and SVs in a single mucosal melanoma. Outer circle depicts canine chromosome number. Blue triangles are SNVs located within coding regions. The middle circle denotes CNVs with gains (in red) and losses (in green) according to the aberration amplitude. Blue or red lines transecting the plot show translocations. (b) Numbers and types of coding mutations identified by SI-WGS and LI-WGS in the discovery cohort. *NDl0-361 and NDl0-363 are independent primary tumors from the same dog. (c) Integrated genomic data is presented for 34 canine melanomas and 3 canine melanoma cell lines. Each column represents data from a single tumor. Indication of tumor type (mucosal, uveal, acral, and cutaneous) is displayed above annotation of recurrently-mutated and hallmark genes. Mutations identified by WGS, aCGH, SNP array, and targeted sequencing are presented in order of frequency as are recurrent CNV regions identified by SNP array and GISTIC as well as recurrent regions involved in translocations identified by LI-WGS. Genomic analysis annotation, tumor ID, and figure legend are presented at the bottom of the figure.

For WGS and LI-WGS respectively a median of 38/11-fold sequence coverage and 209/155-fold physical coverage was achieved **(Supplementary Table 2)**. Read alignment was performed using the canine reference genome CanFam 3.1 and stringent criteria were used to call somatic sequence variants intersecting Seurat, Strelka and Mutect (Materials and Methods). A total of 31,053 somatic single nucleotide variants (SNVs) and small insertions and deletions (indels) were found with a median of 4,223 genome-wide SNVs (range 1,880-6,342) and 316 indels (range 88 - 655) and a median mutation rate of 2.03 mutations per callable haploid megabase (range 0.97-3.14, **Table 2)**. The genome-wide SNV spectrum showed C:G>T:A transitions to be most prevalent, at a median of 27.09% of total SNVs followed by T:A>C:G transitions (median of 21.19%) and C:G>A:T transversions (median 15.74%, **Supplementary Figure 2A**). Despite the prevalence of C:G>T:A transitions, most occurred in CpG dinucleotides and were not enriched at dipyrimidines (median 22.5%). Therefore, a canonical UV signature was not present in any of these cases (**Supplementary Figure 2B**)**(45, 46)**. We additionally looked for *TERT* promoter mutations, which have been reported in 71% of human cutaneous melanomas and are associated with UV damage(47), but no mutations were found within one kilobase of the *TERT* transcription start site. While no single mutation was represented at greater than 4% of the SNV population, C:G>T:A in GCG trinucleotides was the most common mutation (median 6.7%) followed by C>T in ACG (median 2.6%) and C>A in TCT (median 2.5%) (**Supplementary Figure 2C**). No evidence of localized hypermutation (kataegis) was identified in these tumors(48).

**Table 2.**
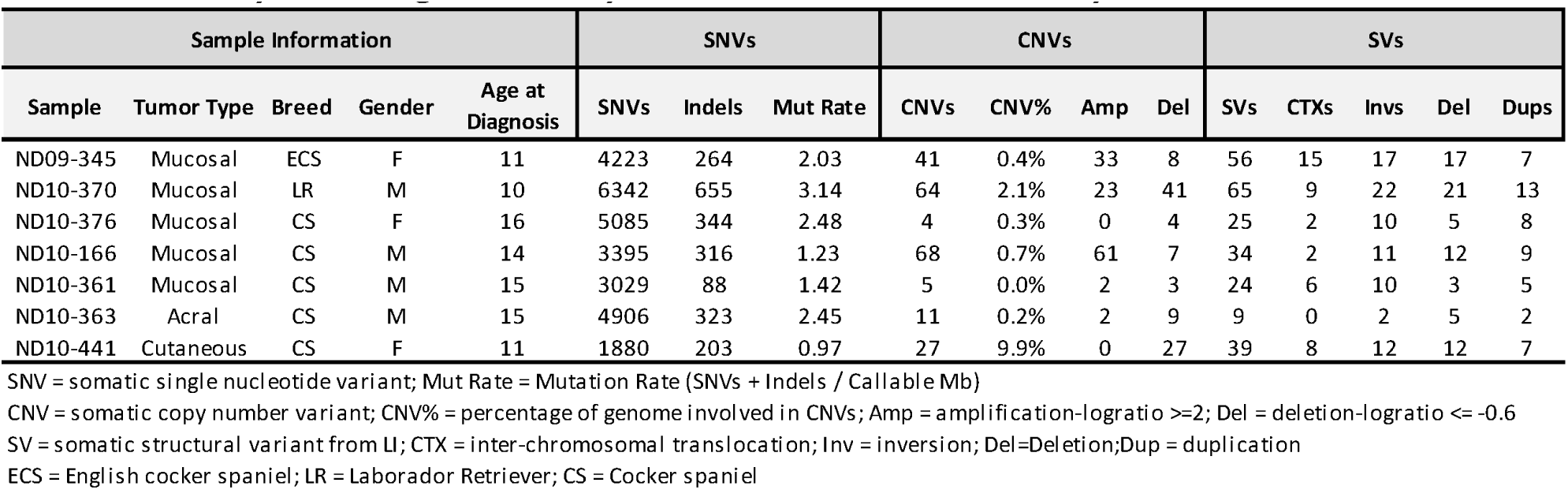
Summary of whole-genome analysis in canine melanoma discovery cohort

**Figure 2.**
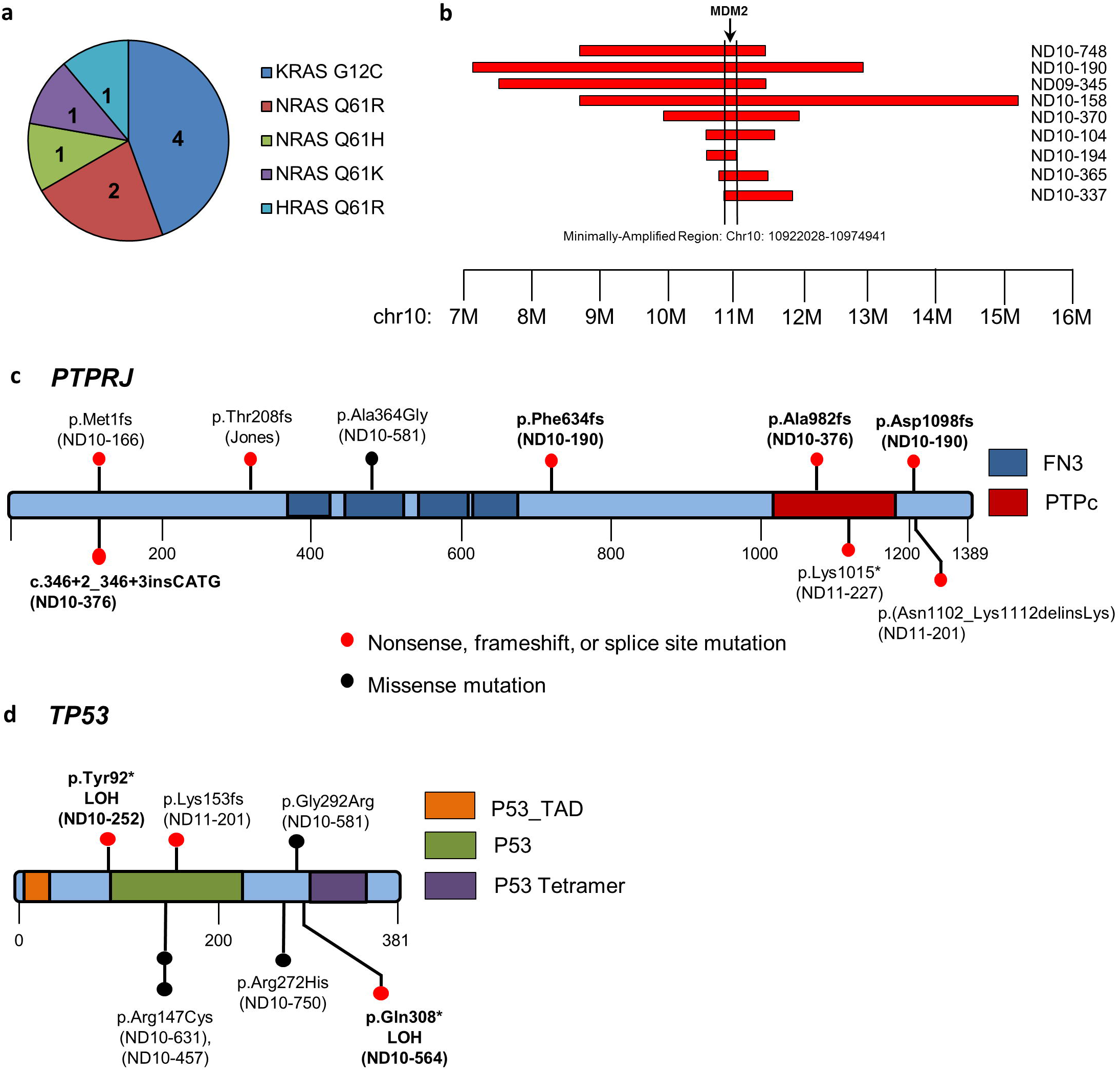
Recurrent somatic alterations in canine melanoma. (a) Distribution of RAS mutations within the cohort of 37 samples (n=9). (b) Recurrently amplified region on CFA 10 found in nine tumors which is defined by the minimal region surrounding *MDM2*. (c) Location of potentially deleterious mutations present in the putative tumor suppressor *PTPRJ* found through Sanger sequencing of the coding sequence for each tumor. (d) Individual mutations and their locations within TP53.

### Somatic coding mutations identified in canine melanoma

Tumors assessed by whole-genome analysis displayed an abundance of somatic SVs and copy number variants (CNVs), with a modest burden of SNVs in coding regions (**Figure 1A and 1B**). The landscape of somatic mutations in the full cohort of 37 tumors based on multi-platform analysis is shown in **Figure 1C**. Circos plots depicting somatic alterations in each tumor in the discovery cohort are shown in **Supplementary Figure 3**. Of the genome-wide SNVs described above, a median of 26 nonsynonymous single-base substitutions and indels occurred within coding regions (nsSNVs, range 14-42) with a median nonsynonymous: synonymous mutation ratio of 2.3 (range 1.9-3.9) (**Figure 1B**). We additionally performed RNA sequencing in this cohort, aligning with TopHat and utilizing IGV to manually validate expressed sequence variants (Materials and Methods). Eighty-five percent of nsSNVs (all but 28) identified by WGS were confirmed by their presence in two or more sequencing platforms **(Supplementary Table 3)**.

**Figure 3.**
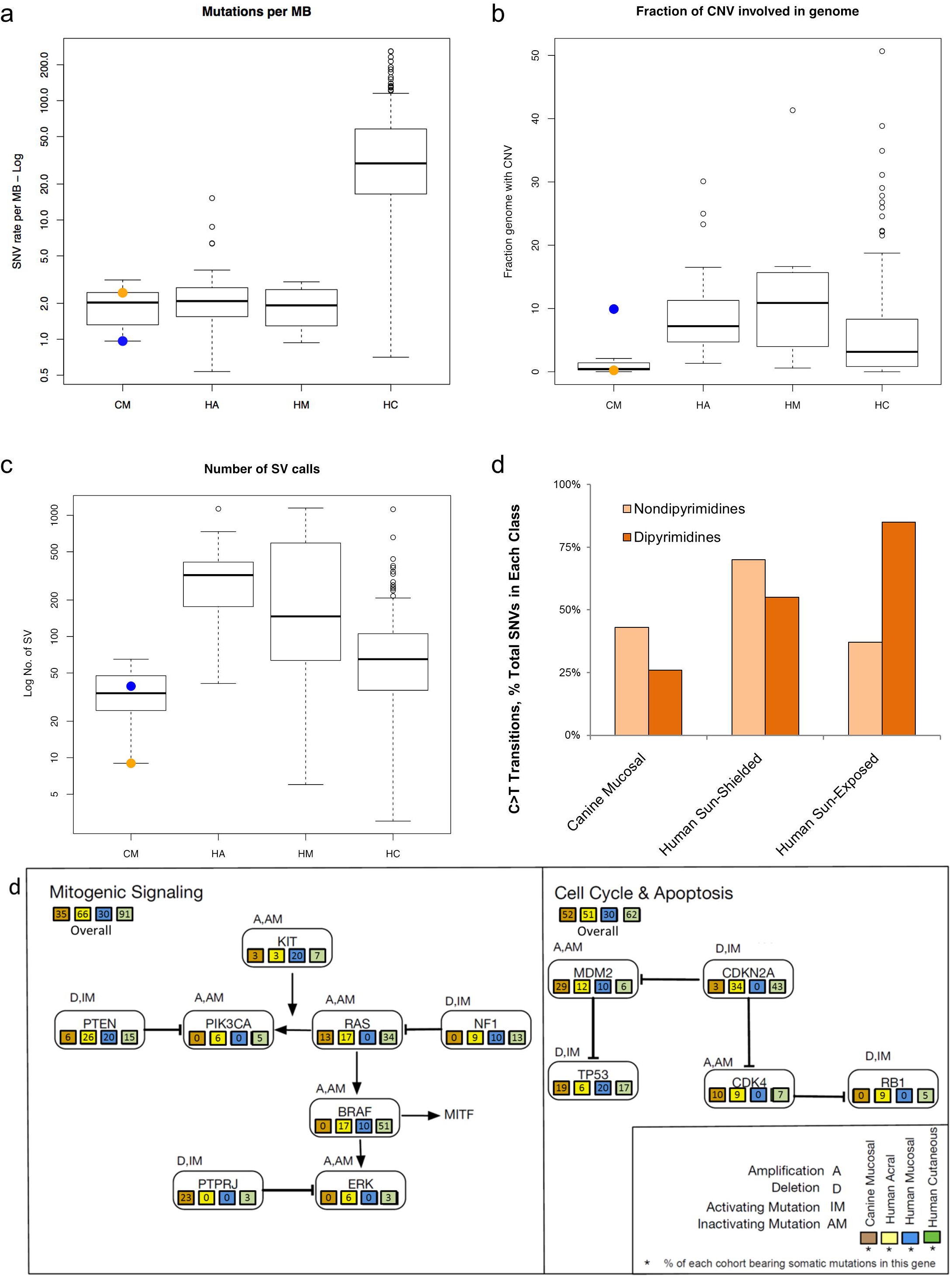
Key dysregulated pathways in canine and human melanoma. (a) Mutation rate in canine and human melanoma subtypes is shown as somatic SNVs per DNA Mb based on WGS in our discovery cohort compared to WGS data from 140 human cutaneous, 35 acral, and 8 mucosal melanomas (Hayward *et al.* 2017). CM: Canine Mucosal, HA: Human Acral, HM: Human Mucosal, and HC: Human Cutaneous Melanoma. Orange and blue dots in the CM plots represent the individual acral and cutaneous subtypes, respectively, in our discovery cohort. (c) Fraction of copy-number-altered genome in canine melanoma and human melanoma sequencing cohorts. (c) Total number of structural variants identified in canine and human melanoma sequencing cohorts. (d) Comparison of C>T transitions in the major melanoma types in dipyrimidine versus non-dipyrimidines. (d) Overall frequency of mutations in key melanoma pathways in our full cohort of 31 mucosal tumors compared to WGS in other subtypes from Hayward *et al.* 2017. Note that, unlike copy number data, sequence data for CDKN2A, ERK, PlK3CA, and RBl were only available for the seven tumors in our discovery cohort.

A number of mutations in orthologs of human cancer genes were present in a single tumor each and include: *ATF6*, *EPAS1*, *FAT2*, *FAT4*, *FOXA3*, *FOXO1*, *GAB2*, *HRAS*, *KIT*, *KRAS*, *MMP21*, *NRAS*, *PBX1,* and *XPO1*. Although no recurrent SNVs were seen in the discovery cohort, three genes were mutated in two cases: *FAT4*, *LRFN2*, and *PTPRJ*. Of these, only *PTRPJ* was validated in multiple platforms in both cases. Both cases containing somatic *PTPRJ* mutations were mucosal (ND10-166 and ND10-376) and both putatively contained two hits. To determine the prevalence of mutations in a panel of genes whose orthologs are known to play a role in human melanomagenesis, as well as the *PTPRJ* gene mutated in two cases, we performed targeted Sanger sequencing of all protein-coding regions of *BAP1, BRAF*, *CDK4*, *GNA11*, *GNAQ*, *KIT*, *KRAS*, *MDM2*, *MITF*, *NF1*, *NRAS*, *PTEN*, *PTPRJ*, and *TP53* in the expanded cohort. *BRAF*, *CDK4*, *GNAQ*, *MDM2*, *MITF,* and *NF1* were all found to be universally wild-type whereas putative pathogenic mutations were discovered in *BAP1, GNA11*, *KIT*, *KRAS*, *NRAS*, *PTEN*, *PTPRJ*, and *TP53* as described below and in **Supplementary Table 4**.

### Somatic copy number and structural variants identified in canine melanoma

Somatic CNVs in the discovery cohort were identified by analysis of short-insert whole genome sequencing (SI-WGS) using established methods (Materials and Methods). A median of 27 focal CNVs (range 4-68), two focal amplifications with a log2 ratio ≥ 2 (range 0-61), and eight focal deletions with a log2 ratio ≤ 0.2 (range 3-41) were identified in the discovery cohort **(Table 2 and Supplementary Table 5)** comprising 0%-10% of the genome (**Table 2**). CNVs were additionally identified in this cohort utilizing Illumina CanineHD BeadChip Single Nucleotide Polymorphism (SNP) arrays and Agilent SurePrint G3 Canine Genome CGH microarrays as previously described(41, 49) (Materials and Methods) with a high platform concordance (**Supplementary Figure 4**). CNV analysis was then expanded to a total of 37 melanomas through SNP arrays in an additional 30 cases in the prevalence cohort (**Table 1 and Supplementary Table 5**). Altered regions were assessed by GISTIC(50) for statistically significant frequency and amplitude (G-score >1.0 and Q<0.05). Ten significant regions were identified including losses within CFA 1, 11, 15, and X, as well as gains in CFA10, 11, 13, 30, and X (**Supplementary Table 6**). Nine of 10 GISTIC regions contained genes and included gains in orthologs of the human cancer genes *MDM2* and *CDK4*. Additional cancer driver alterations (homozygous deletions of tumor suppressor genes or focal amplifications of oncogenes) included *CDKN2A* homozygous deletion (3%) and *KIT* focal amplification (8%) (**Supplementary Table 7**).

Somatic SVs including translocations, inversions, and duplications, were identified in the discovery cohort, based on calls from Delly(51) in LI-WGS (Materials and Methods). Between 9 and 65 predicted SVs were identified in each tumor (median 34) and were predominantly inversions (**Table 2 and Supplementary Table 8**). No recurrent rearrangements were present. Notable alterations in human cancer gene orthologs impacted by SVs in single cases include an *ARHGEF12* inversion, a *BIRC3* inversion, a *CLPTM1L-TERT* translocation, a *DDIT3* inversion, a *MYO5A* translocation, and a *TCF12* inversion. However, two regions of CFA10 and 30 were found to contain somatic SVs in two or more tumors. CFA10 rearrangements occurred in five of seven cases, four of which bore alterations in the region spanning 10 – 12 Mb (also a significant GISTIC region from CNV analysis). CFA30 SVs were also present in three tumors with alterations occurring within a region spanning 15-24 Mb (also encompassing a GISTIC region) in each case. Complex chromosomal rearrangements reminiscent of chromothripsis were observed in four tumors (ND09-345, ND10-370, ND10-361, and ND10-441), with chained or clustered breakpoints localized to a subset of chromosomes in regions that also contained copy-number oscillations(52) (**Supplementary Figure 3**). Gene fusions were also identified in RNAseq data using the TopHat-Fusion software package(53) and IGV verification (Materials and Methods and **Supplementary Table 8**). Three fusions were identified in two tumors (*OSBPL11*-*NFKB1* and *DGKA*-*ABCC5* in ND09-345, and *RPTOR*-*TIMP2* in ND10-376) for which translocations were validated in LI-WGS on IGV inspection. No *BRAF* fusions were identified.

### *BRAF, RAS, NF1*, and *KIT* mutations

Approximately 90% of human cutaneous melanomas are driven in part by *BRAF*, *RAS*, *NF1*, and *KIT* mutations that confer constitutive mitogenic signaling through the MAPK pathway(24, 45, 54). However, these alterations are far less common in human mucosal and acral melanomas(20, 22, 23, 55-57). No somatic alterations in *BRAF* were identified within any platform in our canine melanoma cohort. However, *RAS* family members, whose protein products are predicted to share 100% sequence identity with their human orthologs, were the most commonly mutated genes in aggregate, occurring in 24% of cases in human-conserved hotspots (**Figure 1C and 2A**). *NRAS* codon 61 (Q61R/H/K) and *KRAS* codon 12 (G12C) mutations occurred each in four cases while a single case bore an *HRAS* Q61R mutation (nine total RAS mutations). All three acral and two cutaneous cases bore *NRAS* or *KRAS* mutations, while only 4/31 (13%) of mucosal cases bore an *NRAS*, *KRAS*, or *HRAS* mutation. Although *NF1* copy number losses occurred in six cases, no homozygous deletions or truncating mutations were identified (**Supplementary Table 7**). *KIT* mutations were present in one cutaneous and two mucosal tumors (**Supplementary Tables 3 and 4**). In the cutaneous case, the mutation results in a glutamine (Q) to arginine (R) change in codon 396, notably a site of variation between canine and human orthologs, a change that is not predicted to be damaging by PROVEAN, and may constitute a germline SNP, but germline DNA was not available in this case(58). *KIT* mutations in the mucosal cases included an in-frame deletion of amino acids 560-562, a likely damaging mutation in a commonly mutated region of the human ortholog, as well as an aspartic acid (D) to valine (V) change in codon 815 corresponding to the most common hotspot D816V mutations occurring in the kinase domain of *KIT* in human cancers (**Supplementary Figure 5**)(59). Copy number gains encompassing *KIT* were also present in 10 samples (eight mucosal, one acral, and one cutaneous – Jones, 17CM98, ND10-104, ND10-158, ND10-365, ND10-370, ND10-376, ND10-361, ND10-363, and ND10-441), although no focal amplifications were identified (**Supplementary Table 7**).

### *PTPRJ* Mutations

The most commonly mutated gene in this cohort was the putative tumor suppressor gene *PTPRJ,* not previously shown to have frequent inactivating point mutations in cancer (**Figure 1C and 2C**). PTPRJ (also known as density-enhanced phosphatase 1 (DEP-1) or CD148) is a protein tyrosine phosphatase receptor originally discovered by virtue of its overexpression in dense cultures of human lung fibroblasts(60). It has since been shown to be frequently involved in allelic loss or loss of heterozygosity (LOH) in human cancers and mouse models(61, 62) and to potentially play a role in oncogenesis in diverse cancer types, but somatic homozygous deletions or truncating mutations have yet to be described in cancer from any species and its tumor suppressor status remains controversial(63- 71). Canine and human orthologs share 70% sequence identity with a highly conserved C terminus containing the protein tyrosine phosphatase catalytic domain that is nearly 100% identical between species (**Supplementary Figure 6**). Sequencing of *PTPRJ* across all 37 tumors revealed nine mutations in seven cases (all mucosal), comprising 19% of all tumors and 23% of mucosal cases. Six frameshifts or stop gains were discovered in addition to two splice site mutations, a C-terminal 10-amino acid deletion, and a single predicted damaging missense mutation. Two cases – ND10-190 and ND10-376 – contained two mutations each, consistent with putative bi-allelic inactivation of a tumor suppressor gene. Further, LOH was evident by analysis of adjacent SNPs in WGS data in case ND10-166 bearing the M110fs mutation (**Supplementary Table 10**). Consistent with this finding, the *PTPRJ* frameshift in the ND10-166 tumor occurred at an allele ratio of 61% in DNA and 100% in RNA.

### *MDM2* Amplifications and *TP53* Mutations

Inactivation of the p53 network is a critical step in tumorigenesis in nearly all cancers(72). Both truncating *TP53* mutations and amplifications of *MDM2*, a negative regulator of p53, are key routes to p53 inactivation(73). Although *TP53* mutations and *MDM2* amplifications in human melanoma less common(23-25, 45, 54, 56), 16/37 (43%) of the cases in our cohort of canine melanoma bore focal amplifications of *MDM2* or truncating *TP53* mutations (**Figure 1C**). A recurrent focal amplification on CFA10 was identified by whole genome analysis in three of seven tumors in the discovery cohort with extended SNP array analysis in the prevalence cohort revealing an additional eight tumors bearing these amplifications (minimal region 10.9-11.8 Mb) (**Figure 1C and 2C**). In total, 11/38 cases (29%) bore this amplification involving seven genes, with *MDM2* being the likely amplification target (**Figure 2B**). All such amplifications occurred in mucosal melanomas (11/31, 35%). *CDK4*, a cancer gene 10 Mb proximal to *MDM2* in both human and canine genomes and often the target of bipartite amplification alongside *MDM2(74, 75)*, was co-amplified in three of these cases. Twenty tumors were additionally assessed for MDM2 expression by IHC (**Supplementary Table 11 and Supplementary Figure 7**). Three of five cases with *MDM2* focal amplifications also showed prominent MDM2 staining while no cases lacking *MDM2* amplifications were positive by IHC.

We additionally discovered seven tumors with mutations in *TP53* whose protein product shares 80% identity with its human ortholog (**Supplementary Figure 8**). Three of these mutations were truncating – a homozygous T90X in ND10-252, heterozygous K151fs in ND11-201, and a heterozygous Q306X in ND10-564 (**Figure 2D and Supplementary Table 4**). Of the three missense mutations, R145C and R270H were predicted to be damaging. R145C occurred in two tumors and R270H in a single tumor, with both mutations confirmed somatic through analysis of matched germline DNA. Codon 270 in canine *TP53* is homologous to codon 282 in human *TP53*, the fifth most common hotspot for mutations in human cancer(59). The missense G290R variant is a likely SNP. It occurs in a tumor for which matched germline DNA is unavailable and it is predicted to be neutral, although it has not been previously described(76-78). In keeping with findings in other cancers, no sequence mutations were present in *MDM2* and *MDM2* amplifications were mutually exclusive with *TP53* mutations. Further, *TP53* and *MDM2* alterations were mutually exclusive with *RAS* mutations in all but one case (ND10-748, **Figure 1**).

### Pathway dysregulation in canine melanoma

Common genomic subtypes of human cutaneous melanoma (*BRAF*, *RAS* (N/H/K), and *NF1* in 90% of cases) that engage oncogenic signaling through the MAPK pathway are less common in human non-cutaneous melanoma and in canine malignant melanoma (24% of cases here, **Figure 1C**). Therefore, to undertake unbiased identification of pathways contributing to canine melanomagenesis, we performed pathway analysis using WGS data from the discovery cohort. We generated a list of all genes bearing nonsynonymous mutations, lying within chromosomal breakpoints or significant CNV regions from GISTIC (n=1047) in order to determine enrichment of these mutated genes within specific KEGG and Reactome pathways (Materials and Methods)(79-81). Network analysis of the affected genes identified 97 pathways with significant Benjamini-Hochberg corrected *P*-values (**Supplementary Table 12**). The most significantly enriched pathways were Insulin Receptor Substrate (IRS)-mediated signaling, and IRS-related events, for which 23% (19 genes) of the pathway members are mutated in this cohort. Notably, these pathways converge on MAPK and PI3K mitogenic signaling and contain core pathway members such as *FGF*s, *EIF4G1*, *HRAS*, *KRAS*, *NRAS*, and *RPTOR*. Indeed the majority of the enriched pathways contain members of MAPK, PI3K, or growth factor receptor signaling (e.g. PI3K cascade P=0.002, mTOR signaling P=0.008, signaling by Rho GTPases P=0.012, VEGF signaling P=0.017, RAF activation P=0.017, melanoma signaling P=0.021, RAS signaling P=0.031, and MEK activation P=0.036) and, in many cases, intersections with MDM2 signaling.

## DISCUSSION

Melanoma is a clinically significant disease in dogs, the study of which holds untapped potential for developing clinical approached to improve the lives of pet dogs while also informing human melanoma biology and treatment. Few treatment options are available for locally advanced or metastatic canine melanoma in part because the molecular etiology is still largely unknown. Similarly, limited molecular understanding of rare sun-shielded and *BRAF*wt subtypes of human melanoma has constrained clinical innovation. In order to identify the molecular alterations underlying canine melanoma, we undertook a comprehensive multi-platform genomic investigation. Our integrated analysis confirms that although these tumors are driven by mutational landscapes distinct from those in human cutaneous melanoma, they share important similarities with *BRAF*wt and rare histological subtypes of human melanoma. These data not only guide biological and therapeutic studies in canine melanoma, but they also lend further support for the use of the naturally occurring canine model in comparative studies of human cancers.

This study builds on knowledge of the cytogenetic landscape of canine melanoma(41) to provide a comprehensive view of numbers and types of somatic coding mutations in this cancer. Given the dearth of genomic data for canine melanoma, we initially focused here on collecting primary tumors from diverse breeds. Although numbers were too small to power such analyses, we saw no significant breed-associated alterations in this cohort. Breed-specific somatic mutational landscapes have been shown to occur for other canine cancers such as lymphoma(82). Future expanded study of breed-specific cohorts will be critical for further understanding the role of germline variation in shaping somatic cancer landscapes across species. It will also be important to further define subtype differences in expanded cohorts of canine acral and cutaneous tumors as well as benign and precursor lesions.

Overall, the genomic landscapes of human melanoma vary by anatomic site and degree of sun exposure(22, 26, 57). Cutaneous sun-exposed melanoma is characterized both by high point mutation frequencies linked to UV damage(45) and also only modest burdens of structural variation. In contrast, sun-shielded and non-cutaneous melanomas harbor a low point mutation, but high structural mutation burden. Here, we establish that the canine malignant melanoma genome landscape resembles that reported in human sun-shielded melanoma. Canine melanoma of all subtypes in our discovery cohort is likely sun-shielded, including cutaneous tumors which occur in densely hair-bearing skin, although cropping or shaving during summer months may in some cases increase UV exposure. In keeping with this status, WGS in this cohort provides a deep view of genome-wide mutation burden revealing low point mutation frequencies (median 2.03 somatic mutations per Mb) similar to that seen in human acral and mucosal melanoma WGS data from Hayward *et al.* 2017 (**Figure 3A**)(26). This low point mutation burden relative to human sun-exposed melanoma has potential bearing on expected responses to immunotherapy such as anti-CTLA4 and anti-PD1 checkpoint blockade. Numerous studies have shown a clear positive correlation between mutation burden, abundance of neoantigens, and clinical benefit in human melanoma and other cancers(83, 84). Nonetheless, other molecular determinants of response to immunotherapy exist beyond simply mutation burden and the activity of such agents in canine malignant melanoma remains to be determined. Notably, CNV and SV burden from our WGS in canine malignant melanoma was markedly lower than all subtypes as described in Hayward *et al.* (**Figure 3B and 3C**) (26).

WGS additionally provides a deep view of genome-wide mutation signatures. High point mutation burden in sun-exposed cutaneous melanoma is understood to result from UV-induced over-representation of C>T transitions occurring in dipyrimidines versus non-dipyrimidines. UV-induced C>T mutations occurring in dipyrimidines comprise a low proportion of total SNVs in our cohort (25%), reflective of human sun-shielded cutaneous, mucosal and acral melanoma, in contrast to 85-90% of C>Ts occurring in dipyrimidines in human sun-exposed melanoma (**Figure 3C**)(24, 26, 45, 55, 56, 85). This lends support for a non-UV etiology of canine melanoma.

The genome-wide SNV spectrum further revealed that C>T transitions in CpGs were the most common sequence alterations (**Supplementary Figure 2A**). These mutations correlate with age in human cancers and are due to spontaneous deamination of 5-methylcytosine(46). Enrichment for these mutations in canine melanoma is not surprising given that the largest risk factor for cancer in humans and dogs is biological (not chronological) age(86-91) and that the mean age of these dogs at the time of surgical resection was 13 years (range: 10 – 16). Relative to the average number of human somatic mutations, these data provide further evidence that not only cancer incidence, but also mutational burden increases with biological, rather than chronological, age(92). Commonly observed mutational patterns in human melanoma such as kataegis were not observed, although four tumors exhibited clustered or chained translocations suggestive of breakage-fusion-bridge events due to telomere crisis or of chromothripsis, in which one or a few chromosomes undergo punctuated shattering and reassembly events(52). Such events have been linked to poor outcome in human melanoma(93) and may be enriched in tumors with p53 dysfunction or those that lack means to extend telomeres(94, 95). Notably, we show here that *MDM2* and mutually exclusive *TP53* alterations are common in canine melanoma. Similarly, inactivating p53 mutations have been found in human mucosal and acral melanoma, suggesting p53 pathway dysregulation may be crucial in non-UV induced melanoma development. Further, UV-induced *TERT* promoter mutations are common in human cutaneous melanoma, and, although they are rare in sun-shielded subtypes, these subtypes have been shown to bear enrichment for other types of mutation that drive TERT overexpression such as SVs and CNVs(57). The cutaneous tumors in this cohort do not bear somatic *TERT* promoter mutations or other known genetic lesions that would enable telomere extension. Thus, telomere crisis and the survival of structurally aberrant genomes may play a significant role in the molecular etiology of canine and non-UV induced human melanoma.

Our comprehensive analysis of canine melanoma reveals that most canine melanomas bear a low coding mutation burden and are also less structurally complex than human melanoma. Two WGS approaches coupled with array-based platforms have enabled deep interrogation of these changes, complementing recent cytogenetic analyses of this tumor type(41). Significant copy number gains on CFA10 and 30 that have been reported as a defining signature of these lesions are recapitulated in this dataset (**Supplementary Table 6**). Our multi-platform approach was also able to further elucidate complex chromosomal rearrangements present in these regions. Both regions are involved in multiple intra- and inter-chromosomal structural events across this cohort (**Supplementary Table 8**). Additionally, focal amplification of the CFA10 10-12MB region encompasses *MDM2,* a gene which is known to drive human cancers and is currently being explored as a drug target in TP53 wild type tumors(96). CNVs associated with canine melanoma also include gain of CFA13 and loss of CFA22. While not statistically significant via GISTIC in this cohort, both events are present in individual samples. Overall, extensive copy number and structural variation suggest high levels of large-scale chromosome instability, i.e. gain and loss of whole chromosomes or chromosome arms. Intriguingly, mutually exclusive focal amplification of *MDM2* or inactivating mutation in *TP53* have been shown to be enriched in *BRAF*-, *NRAS*-, and *NF1*-wild-type human melanoma, although human *TP53*-mutant melanomas tend to also display higher mutation burden and presence of C>T transitions(97). Taken together the high degree of structural complexity, the lack of *TERT* mutations or telomere-lengthening mechanisms, and the frequency of *MDM2/TP53* mutations all suggest that chromosome instability plays a key role in canine melanomagenesis.

In the discovery cohort, putatively pathogenic somatic mutations in orthologs of human cancer genes were present in a single tumor each including *ATF6*, *EPAS1*, *FAT2*, *FAT4*, *FOXA3*, *FOXO1*, *GAB2*, *HRAS*, *KIT*, *KRAS*, *MMP21*, *NRAS*, *PBX1,* and *XPO1* (**Supplementary Table 3**). Of the 14 melanoma hallmark genes evaluated in the prevalence cohort (including *PTPRJ*), an additional 24 putatively pathogenic somatic mutations were identified in seven genes – *NRAS*, *TP53*, *PTPRJ*, *KIT*, *KRAS*, *GNA11*, and *BAP1* (**Supplementary Table 4**). Overall, across discovery and prevalence analyses, RAS gene family members were the genes most commonly bearing somatic SNVs, occurring in 24% of cases (**Figure 1C and 2A**), followed by *TP53* and *PTPRJ* mutations each in 19% of cases, *KIT* in 8% and *PTEN* in 5%. Combined, these mutations most commonly impact proliferative and cell cycle/apoptosis pathways in patterns similar to those observed in human melanoma (**Figure 3D**). These findings also suggest that both MAPK pathway inhibition (via MEK inhibitors) or p53 pathway inhibition (via MDM2 inhibitors) may be of equal relevance in canine melanoma as they are in human(38).

The oncogenic MAPK pathway is critically important in many cancers given its central role in conveying extracellular signals to the nucleus in order to regulate cancer hallmarks including proliferation, invasion, metastasis, and angiogenesis. The majority of human cutaneous melanomas are driven in part by constitutive activation of the MAPK pathway through mutation of genes such as *BRAF*, *NRAS*, *NF1*, *KIT*, *GNAQ*, and *GNA11*, often in a mutually exclusive pattern(98). The high frequency of these mutations has motivated the TCGA classification of these tumors according to MAPK mutation status: *BRAF* (∼50% of cases), *RAS* (∼30%), *NF1* (∼15%), and TWT (∼10%)(97). These genomic categories are correlated with clinical, pathological, molecular, and biological features of melanoma and thus may comprise distinct subtypes. However, less common histological subtypes of melanoma such as mucosal, acral, and uveal melanoma bear unique mutation spectra that are not uniformly centered on canonical activating mutations in the MAPK pathway. Correspondingly, it has been shown that *BRAF* mutations are exceedingly rare in predominantly oral canine malignant melanoma and, to date, few alterations in other MAPK members have been discovered. These findings were recapitulated in our cohort, which showed no canonical *BRAF* or *NF1* mutations. Nonetheless, MAPK and/or PI3K signaling have been shown to be activated in nearly all cases(99). Additional mutations impacting the MAPK and PI3K pathways include three *KIT* mutations, two *PTEN* mutations, and one *GNA11* mutation. In total, 35% of mucosal and 43% of all canine melanomas bear an alteration impacting the MAPK pathway (**Figures 1C and 3D**). Prior to our studies described here, the mutations underlying such activation have remained largely unknown.

Here we show a complete absence of somatic *BRAF* mutations (SNVs, CNVs, translocations, or fusions encompassing the *BRAF* locus) in canine malignant melanoma in keeping with prior studies(32, 37, 41, 100). We also did not uncover truncating SNVs in or homozygous deletions of *NF1*. A higher proportion of our cohort bear *RAS* mutations than the 6-13% previously described(32, 99), although prior studies have focused almost exclusively on *NRAS* exons one and two. All three major RAS family members are highly conserved (100% protein identity) between canine and human. In humans, of these family members, malignant melanomas predominantly bear *NRAS* mutations with only very rare *KRAS* and *HRAS* mutations. In our cohort, we found four *NRAS* codon 61 alterations (11%), four *KRAS* G12C mutations and one *HRAS* Q61R mutation. Further, four of these RAS alterations (two *NRAS*, one *KRAS,* and one *HRAS* mutation) occur in mucosal tumors, a frequency of 13% in this subtype. However, in our cohort all three acral tumors and both cutaneous tumors had detectable *RAS* alterations (three *KRAS* and two *NRAS* mutations). This unusual pattern of *RAS* mutation in canine melanoma may reflect important differences in biological, tissue, and species specificities of RAS family members.

These data point to the genomic lesions underlying MAPK and PI3K activation in a substantial proportion of canine melanomas, and to subtle genetic differences in disease subtypes within and across species. Most striking is the discovery of a putative novel tumor suppressor gene, *PTPRJ*, a receptor-type protein tyrosine phosphatase, which has been genetically and functionally implicated in cancer (61, 62), but for which clear genetic mechanisms of inactivation have yet to establish its definitive role as a canonical tumor suppressor gene. *PTPRJ* consists of an extracellular domain with eight fibronectin III motifs, a transmembrane domain, and an intracellular catalytic domain. It was originally cloned from HeLa cells and characterized by overexpression and hyper-activation in dense cultures of fibroblasts, by regulation of contact inhibition, and by its role in regulation of cancer cell proliferation and invasion[60, 101-106]. Early genetic studies of quantitative trait loci for mouse cancer susceptibility with homologous regions in human cancers pointed to recurrent *PTPRJ* deletions, LOH, and missense mutations in small cohorts of colorectal (49%), lung (50%), and breast (78%) carcinomas in addition to a correlation between *PTPRJ* LOH and colorectal cancer progression(61, 62). Additional sequencing studies in larger cohorts have identified nonsynonymous SNPs in the extracellular fibronectin repeats associated with risk of developing thyroid, colorectal, head and neck squamous cell, and esophageal cancers(67, 70, 107-109). More recently, a subclonal K1017N missense mutation in the non-catalytic cytoplasmic domain of PTPRJ was identified in a primary breast tumor with significant enrichment in a brain metastases and patient-derived xenograft(110). PTPRJ substrates that may mediate its tumor suppressive potential include ERK1/2, Akt, various receptor tyrosine kinases, and Src kinases(42, 111- 115). However, *Ptprj* knockout mice have normal development with no cancer predisposition and thus inactivation of this gene does not appear to be sufficient to induce tumorigenesis(65). Across all TCGA studies published to date, the frequency of mutations and/or homozygous deletions appears to be low (400 altered cases), although truncating mutations have been found to comprise 31 of the 257 mutations identified alongside 56 missense mutations predicted to be of medium or high impact (**Supplementary Figure 9 and Supplementary Table 13**)(116, 117). Only 10 mutations are present in the TCGA human cutaneous melanoma data set (a single homozygous deletion and nine missense mutations) with two missense mutations in desmoplastic melanoma and no detectable mutations in uveal melanoma. However, a related receptor-type protein tyrosine phosphatase, *PTPRD,* is thought play a role in regulation of STAT3 signaling and has been frequently implicated as a tumor suppressor in human cancers through inactivating somatic mutation, focal deletion or methylation in glioma, melanoma, neuroblastoma, colorectal, liver, head and neck, and lung cancers(118-121). In human cutaneous melanoma, *PTPRD* is deleted or truncated in 9-12% of cutaneous cases, but has not been determined to occur at high frequency in rare histological subtypes(49, 55, 56, 119, 122).

Here, we present the first report of recurrent somatic truncating mutations in *PTPRJ* in a naturally occurring cancer. We have discovered seven cases (19%) of canine melanomas bearing somatic *PTPRJ* mutations. Canine and human PTPRJ orthologs share 70% sequence identity with a highly conserved C-terminus containing the protein tyrosine phosphatase catalytic domain (**Supplementary Figure 6**). Sequencing of *PTPRJ* across all 38 tumors revealed nine mutations in seven cases (seven mucosal and one uveal) comprising 19% of all tumors and 23% of mucosal cases. Six frameshifts or stop gains were discovered in addition to one splice site mutation, a C-terminal 10-amino acid deletion, and a single predicted damaging missense mutation. Two cases – ND10-190 and ND10-376 – contained two mutations each, consistent with bi-allelic inactivation of a tumor suppressor gene. Further, loss of heterozygosity (LOH) was evident by analysis of adjacent SNPs in WGS data in case ND10-166 bearing the M110fs mutation (**Supplementary Table 10**). Although regional LOH on chromosome 18 was observed by SNP array in three of six cases bearing single mutations in *PTPRJ*, these regions were not observed to directly overlap the coding region of *PTPRJ*. Overall, the enrichment for PTPRJ truncating mutation in canine malignant melanoma bears intriguing implications both for a previously underappreciated role for this gene in human melanoma (e.g. through as-yet understudied roles for hemizygous deletion(123) and/or epigenetic modifications) and for the possibility of unique mechanisms of tumorigenesis across species.

Through deep integrated genomic analysis combining WGS, LI-WGS, RNA sequencing, aCGH, SNP arrays, and targeted Sanger sequencing we have determined that canine melanoma is driven by extensive chromosomal instability and frequent dysregulation of MAPK and cell cycle/apoptosis pathways. In keeping with prior comparative melanoma studies that have incorporated histology, targeted sequencing, and aCGH(32, 36, 38, 41), this work highlights the striking resemblance of canine malignant melanoma to sun-shielded, *BRAF*wt subtypes of human melanoma. Finally, we have additionally discovered a putative novel tumor suppressor that may reflect unique species-specific biology and/or may highlight a tumor suppressive axis more subtly altered and as-yet underappreciated in human melanoma. This work bears immediate relevance for development of improved diagnostic and treatment approaches in canine malignant melanoma and provides further evidence to credential the naturally occurring canine melanoma model for study of relevant genomic subsets of human melanoma.

## MATERIALS AND METHODS

### Clinical samples, histopathology and sample assessment

Tumor samples and whole blood were obtained under institutional review protocols at the Van Andel Research Institute in collaboration with local specialty veterinary clinics. Material was collected at surgery, immediately snap frozen, and preserved in optimal cutting temperature (OCT) compound. Patient matched control DNA was obtained from peripheral blood mononuclear cells. Each resected tumor was evaluated by a board certified pathologist (BD) to estimate tumor content and extent of tissue heterogeneity. Diagnosis of malignant melanoma was histologically confirmed according to criteria defined by the American College of Veterinary Pathologists in addition to criteria recently established by comparative analyses of canine and human melanoma focusing on architecture, pigmentation, and the presence of differentiation markers(32, 99, 124).

### Immunohistochemistry

Two tissue microarrays (TMAs), designated Dog MEL A TMA and Dog MEL B TMA, consisted of 96 individual dogs and 131 tissue samples placed in duplicate and two tissue samples placed in quadruplicate (272 array spots). Multiple tumors from nine dogs were present on the array and multiple samples from varying sites within the same tumor were present for twelve dogs. Additionally, non-melanoma stromal or control normal tissues were included. TMAs were H&E-stained and evaluated via routine immunohistochemical procedures for melanoma cocktail (anti-melan A, anti-melanosome, and anti-tyrosinase), and antibodies to vimentin, MDM2 and p53. Samples scoring positive for MDM2 staining were then confirmed for positive staining with melanoma cocktail and re-evaluated for p53 staining. Positive staining was counted if at least one of the two duplicate samples could be evaluated for both MDM2 and melanoma cocktail on the TMA. Antibodies were purchased from Santa Cruz Biotechnology or Cell Marque. A total of 98 dogs and 189 spots/samples (132 tumors) met these criteria for evaluation for MDM2 protein expression by IHC. Of these 98 dogs, 18 dogs (17%) had melanocytic tumors positive for MDM2 staining in 33 spots/samples (25%). MDM2 staining was predominantly cytoplasmic highest intensity at junction between epithelial and subepithelial (submucosa, dermis). Staining was observed in both malignant pigmented and amelanotic melanoma and benign melanocytomas. Most intense staining (4+ cytoplasmic and nuclear) was observed in a benign cutaneous melanocytoma from a boxer that had additionally a malignant melanoma (negative for MDM2 staining on the array) and multiple cutaneous mast cell tumors.

### Nucleic acid extraction from tumor tissue and blood

Tissue was disrupted and homogenized in Buffer RLT plus (Qiagen AllPrep DNA/RNA Mini Kit), using the Bullet Blender™, Next Advance, and transferred to a microcentrifuge tube containing Buffer RLT plus and 1.6 mm stainless steel beads or 0.9 mm–2.0 mm RNase free stainless steel beads. Blood leukocytes (buffy coat) were isolated from whole blood by centrifugation at room temperature and resuspended in Buffer RLT plus. All samples were homogenized, centrifuged at full speed, and lysates were transferred to Qiagen AllPrep spin columns. Genomic DNA and RNA were then purified following the manufacturer's protocol. DNA was quantified using the Nanodrop spectrophotometer and quality was accessed from 260/280 and 260/230 absorbance ratios. RNA was analyzed on the Agilent Bioanalyzer RNA 6000 Nano Chip to validate RNA integrity (RIN≥7.0).

### Library construction and next generation sequencing

Three μg of genomic DNA from each sample was fragmented to a target size of 300–350 base pairs (bp). Overhangs in the fragmented samples were repaired and adenine bases were ligated on. Diluted paired end Illumina adapters were then ligated onto the A-tailed products. Following ligation, samples were run on a 3% TAE gel to separate products. Ligation products at 300 bp and 350 bp were selected for each sample, isolated from gel punches, and purified. 2× Phusion High-Fidelity PCR Master Mix (Finnzymes; catalog#F-531L) was used to perform PCR to enrich for these products. Enriched PCR products were run on a 2% TAE gel and extracted. Products were quantified using Agilent's High Sensitivity DNA chip (catalog#5067-4626) on the Agilent 2100 Bioanalyzer (catalog#G2939AA).

Long insert whole genome libraries were constructed as previously described(125) with the following modifications: 1100ng inputs were used; following DNA fragmentation, a bead purification was performed at a 1:1.8 (sample volume: bead volume) ratio; a 1% size selection gel was used; and during library enrichment, 10 PCR cycles was used. Libraries were clustered onto Illumina V3 flowcells (San Diego, CA) using Illumina’s TruSeq PE Cluster Kit V3 (cat#PE-401-3001) and sequenced for paired 100bp reads using Illumina’s TruSeq SBS Kit V3 (cat#FC-401-3002, n=3) on the Illumina HiSeq.

10 ng of total RNA was used to generate whole transcriptome libraries for RNA sequencing. Using the Nugen Ovation RNA-Seq System (cat#7100-08), total RNA was used to generate double stranded cDNA, which was amplified using Nugen's SPIA linear amplification process. Amplified cDNA was input into Illumina's TruSeq DNA Sample Preparation Kit – Set A (cat#FC-121-1001) for library preparation. In summary, 1 μg of amplified cDNA was fragmented to a target insert size of 300 bp and end repaired. Samples were then adenylated and indexed paired end adapters were ligated. Ligation products were run on a 2% TAE gel and size selected at 400 bp. Ligation products were isolated from gel punches and purified. Cleaned ligation products were input into PCR to enrich for libraries. PCR products were cleaned and quantified using the Agilent Bioanalyzer.

Tumor and normal libraries were prepared for paired end sequencing as described above. Clusters were generated using Illumina's cBot and HiSeq Paired End Cluster Generation Kits (catalog#PE-401-1001) and sequenced on Illumina's HiSeq 2000 using Illumina's HiSeq Sequencing Kit (catalog#FC-401-1001).

### Next generation sequencing data analysis

BCL to FASTQ file conversion was performed using Illumina's BCL converter tool. Read alignment was performed with BWA (Burrows-Wheeler Aligner)(126) using the canine reference genome CanFam 3.1. Aligned BAMs were indel (insertion/deletion) realigned and recalibrated using GATK(127, 128) and duplicate pairs marked using Picard (http://broadinstitute.github.io/picard/). Variants were called using Strelka(129), Seurat(130) and MuTect(131) and calls were annotated according to dbSNP 139, SNPs on the Illumina CanineHD BeadChip, and SnpEff-3.5(132). Final somatic SNVs were called by at least 2/3 callers. LI-WGS data were utilized for CNV and SV detection. For CNV detection, read depths at every 100 bases across sequenced regions were determined. Next, normalized log2 fold-changes between tumor and normal were calculated and a smoothing window applied. Tumor allele frequencies of known heterozygous germline SNPs were utilized to evaluate potential false positives and correct biases. Finally, the Circular Binary Segmentation algorithm(133) was used to correct log2 fold-changes. For mutation burden metrics, a focal CNV is included if the log2 change is >=|2|. SV detection was performed utilizing Delly(51). A minimum tumor allele ratio of 0.10 and a minimum quality score of 20 is required for an SV to be called.

RNA sequencing data in FASTQ format from the Illumina HiSeq was checked for quality using cycle-by-cycle quality plots and biases, such as GC content. Reads were aligned to the canine reference genome CanFam 3.1 using the TopHat spliced aligner to generate alignment files in BAM format(134). These data were utilized for validation of expressed sequence variants in IGV. Gene fusions were also identified in RNAseq data using the TopHat-Fusion software package(53) and IGV verification. Results were reported in tables showing p-values (adjusted for multiple testing) and normalized abundance data in terms of FPKM (fragments per kilo-base of transcript per million mapped reads) which were also examined manually. Gene and transcript annotations were downloaded from ENSEMBL (CanFam 3.1.68) and germline SNPs filtered out using publicly available canine SNP data (71- 73).

### Data access

Next generation sequencing data from this study have been submitted to the NCBI Biosample Database (http://www.ncbi.nlm.nih.gov/biosample/7196161) under study ID SUB2752127 and accession numbers SAMN07196161, SAMN07196162, SAMN07196163, SAMN07196164, SAMN07196165, SAMN07196166, and SAMN07196167.

### Pathway analysis

A list of 1,405 genes with single nucleotide variation or structural variation or copy number variation from the discovery cohort were analyzed using ClueGo4(79), a Cytoscape plug-in, to create a functionally organized pathway network. Kappa scores were then used to measure association between the networks. Functional networks were created with a minimum Kappa score threshold of 0.5 and a minimum of 3 affected genes in every network forming at least 10% of the total associated genes in that particular network. The genes were assigned to the networks based on the predefined pathways from KEGG, REACTOME and Wiki Pathways. 97 pathways were obtained, all with Benjamini-Hochberg corrected P-value <0.05. These pathways were grouped together based on inter-term kappa score and named by the most significant pathway in the respective groups.

### PCR amplification and Sanger sequencing analysis

PCR amplification of 15 genes (*NRAS, KRAS, BRAF, GNA11, GNAQ, PTPRJ, TP53, MDM2, BAP1, CDK4, PTEN, c-KIT, MITF* and *NF1*) was performed using primers targeting all coding exons as shown in Supplementary Table 9. All amplification reactions were performed using Platinum Taq DNA Polymerase #10966-034 (Life Technologies; Carlsbad, CA). Briefly, each primer pair was mixed with 10 ng of genomic DNA and subjected to the following cycling parameters: 94°C for 2 min., 3 cycles at each temperature: 30 sec. at 94°C, 30 sec. at 60-57°C, 45 sec. at 72°C; 25 cycles: 30 sec. at 94°C, 30 sec. at 62°C, 45 sec. at 72°C; final extension of 5 min. at 72°C. PCR amplicons were sequenced using M13 forward and reverse primers at the Arizona State University’s DNA Lab (Tempe, AZ).

### Array comparative genomic hybridization

Oligo array CGH (aCGH) was performed by co-hybridization of tumor (test) DNA and a common reference DNA sample, where the latter comprised an equimolar pool of genomic DNA samples from multiple healthy individuals of various breeds. DNA was labeled using an Agilent SureTag Labeling Kit (Agilent Technologies, Santa Clara, CA) with all test samples labeled with Cyanine-3-dCTP and the common reference sample labeled with Cyanine-5-dCTP. Fluorochrome incorporation and final probe concentrations were determined using routine spectrophotometric parameters with readings taken from a Nanodrop1000. Fluorescently labeled test and reference samples were co-hybridized to Canine G3 180,000 feature CGH arrays (Agilent, AMADID 025522) for 40 h at 65 °C and 20 rpm, as described previously(135, 136). Arrays were scanned at 3 μm using a high-resolution microarray scanner (Agilent,Model G2505C) and data extracted using Feature Extraction (v10.9) software. Scan data were assessed for quality by the ‘Quality Metrics’ report in Agilent’s Feature extraction software (v10.5) (Agilent Technologies).

### SNP array genotyping

SNP genotyping was performed using the Illumina CanineHD array (cat#WG-440-1001). Per manufacturer’s protocol, 200ng of DNA was first denatured then neutralized with 0.1N NaOH before amplification at 37°C for 24 hours. The amplified DNA was then enzymatically fragmented and precipitated using 100% 2-propanol before drying for one hour at room temperature. After resuspension the fragmented DNA was then denatured and loaded onto the CanineHD BeadChip and hybridized for 16 hours at 48°C. BeadChips were washed, a single base extension of hybridized primers added followed by multi-layer staining of the primers. Arrays were then washed, coated with the XC4 reagent (Illumina) and dried under vacuum for one hour. Coated arrays were read on the HiScan system and data visualized using the Illumina Genome Studio Genotyping 2.0 software with an average sample call rate of 97%.

### aCGH and SNP array data analysis

For both aCGH and SNP arrays, copy number data were analyzed with NEXUS Copy Number v8.0 software (Biodiscovery Inc., CA, USA). For cross-platform comparisons, LI-WGS BAMs were also analyzed utilizing Nexus software. CNVs were identified using a FASST2 segmentation algorithm with a significance threshold of 5.5×10^-6.^ Aberrations were defined as a minimum of three consecutive probes with log2 tumor: reference value of >1.14 (high gain), 1.13 to 0.2 (gain), -0.23 to -1.1 (loss), <-1.1 (big loss). Recurrent CNVs within each subtype were determined within NEXUS using an involvement threshold of 50 %. Significance of these regions was then determined in NEXUS using the GISTIC algorithm (to identify regions with a statistically high frequency of CNVs over background) with a Gscore cut off of G>1.0 and a significance of Q<0.05. CNV frequency comparisons amongst sample groups were performed in NEXUS using Fisher’s exact test with differential threshold of >50 % and significance p<0.05. Significance of each probe between the two groups was calculated in NEXUS using a Mann– Whitney test for median comparison.

## Acknowledgements

We thank Antonia Pritchard for valued input on genomic analyses. These studies were supported by Brooke’s Blossoming Hope for Childhood Cancer Foundation, The Stand Up To Cancer – Melanoma Research Alliance/Melanoma Dream Team Translational Cancer Research Grant (SU2C-AACR-DT0612, Stand Up To Cancer is a program of the Entertainment Industry Foundation administered by the American Association for Cancer Research), Dell Inc. through the Dell Powering the Possible Program, NIH Grants UM1 CA186689 and RC2 CA148149, and philanthropic support to the TGen Foundation. CGH analysis was supported by the NC State Cancer Genomics Fund (MB).

## References

1. Siegel R, Ma J, Zou Z, Jemal A. Cancer statistics, 2014. CA: A Cancer Journal for Clinicians. 2014;64(1):9–29.

2. Pollock PM, Harper UL, Hansen KS, Yudt LM, Stark M, Robbins CM, et al. High frequency of BRAF mutations in nevi. Nature genetics. 2002;33(1):19–20.

3. Chapman PB, Hauschild A, Robert C, Haanen JB, Ascierto P, Larkin J, et al. Improved survival with vemurafenib in melanoma with BRAF V600E mutation. New England Journal of Medicine. 2011 2011;364(26):2507–16.

4. Davies H, Bignell GR, Cox C, Stephens P, Edkins S, Clegg S, et al. Mutations of the BRAF gene in human cancer. Nature. 2002;417(6892):949–54.

5. Sun C, Wang L, Huang S, Heynen GJ, Prahallad A, Robert C, et al. Reversible and adaptive resistance to BRAF (V600E) inhibition in melanoma. Nature. 2014;508(7494):118–22.

6. Flaherty KT, Robert C, Hersey P, Nathan P, Garbe C, Milhem M, et al. Improved survival with MEK inhibition in BRAF-mutated melanoma. New England Journal of Medicine. 2012;367(2):107–14.

7. Flaherty KT, Infante JR, Daud A, Gonzalez R, Kefford RF, Sosman J, et al. Combined BRAF and MEK inhibition in melanoma with BRAF V600 mutations. New England Journal of Medicine. 2012;367(18):1694–703.

8. Hodi FS, O ’Day SJ, McDermott DF, Weber RW, Sosman JA, Haanen JB, et al. Improved survival with ipilimumab in patients with metastatic melanoma. New England Journal of Medicine. 2010;363(8):711–23.

9. Thakur MD, Salangsang F, Landman AS, Sellers WR, Pryer NK, Levesque MP, et al. Modelling vemurafenib resistance in melanoma reveals a strategy to forestall drug resistance. Nature. 2013;494(7436):251–5.

10. Khanna C, Fan TM, Gorlick R, Helman LJ, Kleinerman ES, Adamson PC, et al. Towards a Drug Development Path that Targets Metastatic Progression in Osteosarcoma. Clinical Cancer Research. 2014:clincanres. 2574.013.

11. Paoloni M, Khanna C. Translation of new cancer treatments from pet dogs to humans. Nature Reviews Cancer. 2008 2008;8(2):147–56.

12. Tang J, Li Y, Lyon K, Camps J, Dalton S, Ried T, et al. Cancer driver–passenger distinction via sporadic human and dog cancer comparison: a proof-of-principle study with colorectal cancer. Oncogene. 2014;33(7):814–22.

13. Liu D, Xiong H, Ellis AE, Northrup NC, Rodriguez CO, O ’Regan RM, et al. Molecular homology and difference between spontaneous canine mammary cancer and human breast cancer. Cancer research. 2014;74(18):5045–56.

14. Bushell KR, Kim Y, Chan FC, Ben-Neriah S, Jenks A, Alcaide M, et al. Genetic inactivation of TRAF3 in canine and human B-cell lymphoma. Blood. 2015;125(6):999–1005.

15. Schiffman JD, Breen M. Comparative oncology: what dogs and other species can teach us about humans with cancer. Phil Trans R Soc B. 2015;370(1673):20140231.

16. LeBlanc AK, Breen M, Choyke P, Dewhirst M, Fan TM, Gustafson DL, et al. Perspectives from man’s best friend: National Academy of Medicine’s Workshop on Comparative Oncology. Sci Transl Med. 2016;8(324):324ps5.

17. Manolidis S, Donald PJ. Malignant mucosal melanoma of the head and neck. Cancer. 1997;80(8):1373–86.

18. Meleti M, Leemans CR, de Bree R, Vescovi P, Sesenna E, van der Waal I. Head and neck mucosal melanoma: experience with 42 patients, with emphasis on the role of postoperative radiotherapy. Head & Neck. 2008;30(12):1543–51.

19. Tanaka T, Yamada R, Tanaka M, Shimizu K, Oka H, editors. A study on the image diagnosis of melanoma. Engineering in Medicine and Biology Society, 2004 IEMBS’04 26th Annual International Conference of the IEEE; 2004: IEEE.

20. Curtin JA, Busam K, Pinkel D, Bastian BC. Somatic activation of KIT in distinct subtypes of melanoma. Journal of Clinical Oncology. 2006;24(26):4340–6.

21. Maldonado JL, Fridlyand J, Patel H, Jain AN, Busam K, Kageshita T, et al. Determinants of BRAF mutations in primary melanomas. Journal of the National Cancer Institute. 2003;95(24):1878–90.

22. Curtin JA, Fridlyand J, Kageshita T, Patel HN, Busam KJ, Kutzner H, et al. Distinct sets of genetic alterations in melanoma. New England Journal of Medicine. 2005;353(20):2135–47.

23. Turajlic S, Furney SJ, Lambros MB, Mitsopoulos C, Kozarewa I, Geyer FC, et al. Whole genome sequencing of matched primary and metastatic acral melanomas. Genome Res. 2012 2012/02/01/;22(2):196–207.

24. Krauthammer M, Kong Y, Ha BH, Evans P, Bacchiocchi A, McCusker JP, et al. Exome sequencing identifies recurrent somatic RAC1 mutations in melanoma. Nature genetics. 2012;44(9):1006–14.

25. Furney SJ, Turajlic S, Stamp G, Nohadani M, Carlisle A, Thomas JM, et al. Genome sequencing of mucosal melanomas reveals that they are driven by distinct mechanisms from cutaneous melanoma. The Journal of Pathology. 2013;230(3):261–9.

26. Hayward NK, Wilmott JS, Waddell N, Johansson PA, Field MA, Nones K, et al. Whole-genome landscapes of major melanoma subtypes. Nature. 2017;545(7653):175–80.

27. Chang AE, Karnell LH, Menck HR. The National Cancer Data Base report on cutaneous and noncutaneous melanoma. Cancer. 1998;83(8):1664–78.

28. Cotchin E. Melanotic tumours of dogs. Journal of Comparative Pathology and Therapeutics. 1955;65:115-IN14.

29. Smith SH, Goldschmidt MH, McManus PM. A comparative review of melanocytic neoplasms. Veterinary Pathology Online. 2002 2002;39(6):651–78.

30. Villamil JA, Henry CJ, Bryan JN, Ellersieck M, Schultz L, Tyler JW, et al. Identification of the most common cutaneous neoplasms in dogs and evaluation of breed and age distributions for selected neoplasms. Journal of the American Veterinary Medical Association. 2011;239(7):960–5.

31. Bergman PJ. Canine Oral Melanoma. Clinical Techniques in Small Animal Practice. 2007 2007/05//;22(2):55–60.

32. Gillard M, Cadieu E, De Brito C, Abadie J, Vergier B, Devauchelle P, et al. Naturally occurring melanomas in dogs as models for non-UV pathways of human melanomas. Pigment Cell & Melanoma Research. 2014;27(1):90–102.

33. Prasad ML, Patel SG, Huvos AG, Shah JP, Busam KJ. Primary mucosal melanoma of the head and neck. Cancer. 2004;100(8):1657–64.

34. Bergman PJ, Wolchok JD. Of mice and men (and dogs): development of a xenogeneic DNA vaccine for canine oral malignant melanoma. Cancer Ther. 2008;6:817–26.

35. Bergman P, Kent M, Farese J. Melanoma. Withrow and MacEwen’s Small Animal Clinical Oncology SJ, Withrow, DM, Vail, and RL, Page, eds(St Louis, MO: Elsevier/Saunders). 2013:321-34. 36.

36. Simpson RM, Bastian BC, Michael HT, Webster JD, Prasad ML, Conway CM, et al. Sporadic naturally occurring melanoma in dogs as a preclinical model for human melanoma. Pigment Cell & Melanoma Research. 2014;27(1):37–47.

37. Shelly S, Chien MB, Yip B, Kent MS, Theon AP, McCallan JL, et al. Exon 15 BRAF mutations are uncommon in canine oral malignant melanomas. Mammalian Genome. 2005;16(3):211–7.

38. Fowles JS, Denton CL, Gustafson DL. Comparative analysis of MAPK and PI3K/AKT pathway activation and inhibition in human and canine melanoma. Veterinary and Comparative Oncology. 2013 2013:n/a-n/a.

39. Murakami A, Mori T, Sakai H, Murakami M, Yanai T, Hoshino Y, et al. Analysis of KIT expression and KIT exon 11 mutations in canine oral malignant melanomas. Veterinary and Comparative Oncology. 2011;9(3):219–24.

40. Chu P-Y, Pan S-L, Liu C-H, Lee J, Yeh L-S, Liao AT. KIT gene exon 11 mutations in canine malignant melanoma. The Veterinary Journal. 2013;196(2):226–30.

41. Poorman K, Borst L, Moroff S, Roy S, Labelle P, Motsinger-Reif A, et al. Comparative cytogenetic characterization of primary canine melanocytic lesions using array CGH and fluorescence in situ hybridization. Chromosome Res. 2014:1–16.

42. Spring K, Lapointe L, Caron C, Langlois S, Royal I. Phosphorylation of DEP-1/PTPRJ on threonine 1318 regulates Src activation and endothelial cell permeability induced by vascular endothelial growth factor. Cellular signalling. 2014;26(6):1283–93.

43. Liang WS, Aldrich J, Tembe W, Kurdoglu A, Cherni I, Phillips L, et al. Long insert whole genome sequencing for copy number variant and translocation detection. Nucleic acids research. 2014;42(2):e8– e.

44. Smedley R, Lamoureux J, Sledge D, Kiupel M. Immunohistochemical diagnosis of canine oral amelanotic melanocytic neoplasms. Veterinary Pathology Online. 2011;48(1):32–40.

45. Berger MF, Hodis E, Heffernan TP, Deribe YL, Lawrence MS, Protopopov A, et al. Melanoma genome sequencing reveals frequent PREX2 mutations. Nature. 2012;485(7399):502–6.

46. Alexandrov LB, Nik-Zainal S, Wedge DC, Aparicio SA, Behjati S, Biankin AV, et al. Signatures of mutational processes in human cancer. Nature. 2013.

47. Huang FW, Hodis E, Xu MJ, Kryukov GV, Chin L, Garraway LA. Highly recurrent TERT promoter mutations in human melanoma. Science. 2013;339(6122):957–9.

48. Nik-Zainal S, Alexandrov LB, Wedge DC, Van Loo P, Greenman CD, Raine K, et al. Mutational processes molding the genomes of 21 breast cancers. Cell. 2012;149(5):979–93.

49. Stark M, Hayward N. Genome-wide loss of heterozygosity and copy number analysis in melanoma using high-density single-nucleotide polymorphism arrays. Cancer research. 2007;67(6):2632–42.

50. Mermel CH, Schumacher SE, Hill B, Meyerson ML, Beroukhim R, Getz G. GISTIC2. 0 facilitates sensitive and confident localization of the targets of focal somatic copy-number alteration in human cancers. Genome Biol. 2011;12(4):R41.

51. Rausch T, Zichner T, Schlattl A, Stütz AM, Benes V, Korbel JO. DELLY: structural variant discovery by integrated paired-end and split-read analysis. Bioinformatics. 2012;28(18):i333–i9.

52. Stephens PJ, Greenman CD, Fu B, Yang F, Bignell GR, Mudie LJ, et al. Massive genomic rearrangement acquired in a single catastrophic event during cancer development. Cell. 2011;144(1):27–40.

53. Kim D, Salzberg SL. TopHat-Fusion: an algorithm for discovery of novel fusion transcripts. Genome Biol. 2011;12(8):R72.

54. Hodis E, Watson IR, Kryukov GV, Arold ST, Imielinski M, Theurillat J-P, et al. A landscape of driver mutations in melanoma. Cell. 2012;150(2):251–63.

55. Furney SJ, Turajlic S, Stamp G, Nohadani M, Carlisle A, Thomas JM, et al. Genome sequencing of mucosal melanomas reveals that they are driven by distinct mechanisms from cutaneous melanoma. The Journal of Pathology. 2013 2013;230(3):261–9.

56. Furney SJ, Turajlic S, Stamp G, Thomas JM, Hayes A, Strauss D, et al. The mutational burden of acral melanoma revealed by whole-genome sequencing and comparative analysis. Pigment Cell & Melanoma Research. 2014;27(5):835–8.

57. Liang WS, Hendricks W, Kiefer J, Schmidt J, Sekar S, Carpten J, et al. Integrated genomic analyses reveal frequent TERT aberrations in acral melanoma. Genome Res. 2017;27(4):524–32.

58. Choi Y, Sims GE, Murphy S, Miller JR, Chan AP. Predicting the functional effect of amino acid substitutions and indels. 2012.

59. Forbes SA, Beare D, Gunasekaran P, Leung K, Bindal N, Boutselakis H, et al. COSMIC: exploring the world’s knowledge of somatic mutations in human cancer. Nucleic acids research. 2015;43(D1):D805–D11.

60. Ostman A, Yang Q, Tonks NK. Expression of DEP-1, a receptor-like protein-tyrosine-phosphatase, is enhanced with increasing cell density. Proceedings of the National Academy of Sciences. 1994;91(21):9680–4.

61. Ruivenkamp CA, van Wezel T, Zanon C, Stassen AP, Vlcek C, Csikós T, et al. Ptprj is a candidate for the mouse colon-cancer susceptibility locus Scc1 and is frequently deleted in human cancers. Nature genetics. 2002;31(3):295–300.

62. Ruivenkamp C, Hermsen M, Postma C, Klous A, Baak J, Meijer G, et al. LOH of PTPRJ occurs early in colorectal cancer and is associated with chromosomal loss of 18q12–21. Oncogene. 2003;22(22):3472–4.

63. Lesueur F, Pharoah PD, Laing S, Ahmed S, Jordan C, Smith PL, et al. Allelic association of the human homologue of the mouse modifier Ptprj with breast cancer. Human molecular genetics. 2005;14(16):2349–56.

64. Godfrey R, Arora D, Bauer R, Stopp S, Müller JP, Heinrich T, et al. Cell transformation by FLT3 ITD in acute myeloid leukemia involves oxidative inactivation of the tumor suppressor protein-tyrosine phosphatase DEP-1/PTPRJ. Blood. 2012;119(19):4499–511.

65. Trapasso F, Drusco A, Costinean S, Alder H, Aqeilan RI, Iuliano R, et al. Genetic ablation of Ptprj, a mouse cancer susceptibility gene, results in normal growth and development and does not predispose to spontaneous tumorigenesis. DNA and cell biology. 2006;25(6):376–82.

66. Petermann A, Haase D, Wetzel A, Balavenkatraman KK, Tenev T, Gührs KH, et al. Loss of the Protein-Tyrosine Phosphatase DEP-1/PTPRJ Drives Meningioma Cell Motility. Brain Pathology. 2011;21(4):405–18.

67. Mita Y, Yasuda Y, Sakai A, Yamamoto H, Toyooka S, Gunduz M, et al. Missense polymorphisms of PTPRJ and PTPN13 genes affect susceptibility to a variety of human cancers. Journal of cancer research and clinical oncology. 2010;136(2):249–59.

68. Iuliano R, Palmieri D, He H, Iervolino A, Borbone E, Pallante P, et al. Role of PTPRJ genotype in papillary thyroid carcinoma risk. Endocrine-related cancer. 2010;17(4):1001–6.

69. Gaudio E, Costinean S, Raso C, Mangone G, Paduano F, Zanesi N, et al. Tumor suppressor activity of PTPRJ, a receptor-type protein tyrosine phosphatase, in human melanoma cells. Cancer research. 2008;68(9 Supplement):131-.

70. Iuliano R, Le Pera I, Cristofaro C, Baudi F, Arturi F, Pallante P, et al. The tyrosine phosphatase PTPRJ/DEP-1 genotype affects thyroid carcinogenesis. Oncogene. 2004;23(52):8432–8.

71. Toland AE, Rozek LS, Presswala S, Rennert G, Gruber SB. PTPRJ haplotypes and colorectal cancer risk. Cancer Epidemiology Biomarkers & Prevention. 2008;17(10):2782–5.

72. Vogelstein B, Lane D, Levine AJ. Surfing the p53 network. Nature. 2000 2000;408(6810):307-10. 73.

73. Momand J, Zambetti GP, Olson DC, George D, Levine AJ. The mdm-2 oncogene product forms a complex with the p53 protein and inhibits p53-mediated transactivation. Cell. 1992;69(7):1237–45.

74. Wikman H, Nymark P, Väyrynen A, Jarmalaite S, Kallioniemi A, Salmenkivi K, et al. CDK4 is a probable target gene in a novel amplicon at 12q13. 3–q14. 1 in lung cancer. Genes, Chromosomes and Cancer. 2005;42(2):193–9.

75. Reifenberger G, Ichimura K, Reifenberger J, Elkahloun AG, Meltzer PS, Collins VP. Refined mapping of 12q13–q15 amplicons in human malignant gliomas suggests CDK4/SAS and MDM2 as independent amplification targets. Cancer research. 1996;56(22):5141–5.

76. Vaysse A, Ratnakumar A, Derrien T, Axelsson E, Rosengren Pielberg G, Sigurdsson S, et al. Identification of Genomic Regions Associated with Phenotypic Variation between Dog Breeds using Selection Mapping. PLoS Genet. 2011 2011/10/13/;7(10).

77. Axelsson E, Ratnakumar A, Arendt M-L, Maqbool K, Webster MT, Perloski M, et al. The genomic signature of dog domestication reveals adaptation to a starch-rich diet. Nature. 2013;495(7441):360–4.

78. Lindblad-Toh K, Wade CM, Mikkelsen TS, Karlsson EK, Jaffe DB, Kamal M, et al. Genome sequence, comparative analysis and haplotype structure of the domestic dog. Nature. 2005 2005;438(7069):803–19.

79. Bindea G, Mlecnik B, Hackl H, Charoentong P, Tosolini M, Kirilovsky A, et al. ClueGO: a Cytoscape plug-in to decipher functionally grouped gene ontology and pathway annotation networks. Bioinformatics. 25(8):1091–3.

80. Kanehisa M, Goto S, Sato Y, Furumichi M, Tanabe M. KEGG for integration and interpretation of large-scale molecular data sets. Nucleic acids research. 2011:gkr988.

81. Croft D, Mundo AF, Haw R, Milacic M, Weiser J, Wu G, et al. The Reactome pathway knowledgebase. Nucleic acids research. 2014;42(D1):D472–D7.

82. Elvers I, Turner-Maier J, Swofford R, Koltookian M, Johnson J, Stewart C, et al. Exome sequencing of lymphomas from three dog breeds reveals somatic mutation patterns reflecting genetic background. Genome Research. 2015;25(11):1634–45.

83. Snyder A, Makarov V, Merghoub T, Yuan J, Zaretsky JM, Desrichard A, et al. Genetic basis for clinical response to CTLA-4 blockade in melanoma. New England Journal of Medicine. 2014;371(23):2189–99.

84. Postow MA, Callahan MK, Wolchok JD. Immune checkpoint blockade in cancer therapy. Journal of Clinical Oncology. 2015;33(17):1974–82.

85. Pleasance ED, Cheetham RK, Stephens PJ, McBride DJ, Humphray SJ, Greenman CD, et al. A comprehensive catalogue of somatic mutations from a human cancer genome. Nature. 2010 2010/01/14/;463(7278):191–6.

86. Tomasetti C, Vogelstein B, Parmigiani G. Half or more of the somatic mutations in cancers of self-renewing tissues originate prior to tumor initiation. Proceedings of the National Academy of Sciences. 2013;110(6):1999–2004.

87. Cohen D, Reif JS, Brodey RS, Keiser H. Epidemiological analysis of the most prevalent sites and types of canine neoplasia observed in a veterinary hospital. Cancer Research. 1974;34(11):2859–68.

88. Dorn CR, Taylor DON, Schneider R, Hibbard HH, Klauber MR. Survey of animal neoplasms in Alameda and Contra Costa Counties, California. II. Cancer morbidity in dogs and cats from Alameda County. Journal of the National Cancer Institute. 1968 1968;40(2):307–18.

89. Dobson JM, Samuel S, Milstein H, Rogers K, Wood JLN. Canine neoplasia in the UK: estimates of incidence rates from a population of insured dogs. Journal of small animal practice. 2002 2002;43(6):240–6.

90. Merlo DF, Rossi L, Pellegrino C, Ceppi M, Cardellino U, Capurro C, et al. Cancer incidence in pet dogs: findings of the Animal Tumor Registry of Genoa, Italy. J Vet Intern Med. 2008 2008/08//Julundefined;22(4):976–84.

91. Albert RE, Benjamin SA, Shukla R. Life span and cancer mortality in the beagle dog and humans. Mechanisms of ageing and development. 1994;74(3):149–59.

92. Turker MS. Somatic cell mutations: can they provide a link between aging and cancer? Mechanisms of ageing and development. 2000;117(1):1–19.

93. Hirsch D, Kemmerling R, Davis S, Camps J, Meltzer PS, Ried T, et al. Chromothripsis and focal copy number alterations determine poor outcome in malignant melanoma. Cancer Research. 2013;73(5):1454–60.

94. Rausch T, Jones DT, Zapatka M, Stütz AM, Zichner T, Weischenfeldt J, et al. Genome sequencing of pediatric medulloblastoma links catastrophic DNA rearrangements with TP53 mutations. Cell. 2012;148(1):59–71.

95. Maher CA, Wilson RK. Chromothripsis and human disease: piecing together the shattering process. Cell. 2012;148(1):29–32.

96. Vassilev LT, Vu BT, Graves B, Carvajal D, Podlaski F, Filipovic Z, et al. In vivo activation of the p53 pathway by small-molecule antagonists of MDM2. Science. 2004 2004;303(5659):844–8.

97. Network CGA. Genomic classification of cutaneous melanoma. Cell. 2015;161(7):1681–96.

98. Zhang T, Dutton-Regester K, Brown KM, Hayward NK. The genomic landscape of cutaneous melanoma. Pigment Cell & Melanoma Research. 2016.

99. Simpson RM, Bastian BC, Michael HT, Webster JD, Prasad ML, Conway CM, et al. Sporadic naturally occurring melanoma in dogs as a preclinical model for human melanoma. Pigment Cell & Melanoma Research. 2013 2013/10//:n/a-n/a.

100. Mochizuki H, Kennedy K, Shapiro SG, Breen M. BRAF Mutations in Canine Cancers. PLoS ONE. 2015;10(6):e0129534.

101. Borges LG, Seifert RA, Grant FJ, Hart CE, Disteche CM, Edelhoff S, et al. Cloning and characterization of rat density-enhanced phosphatase-1, a protein tyrosine phosphatase expressed by vascular cells. Circulation research. 1996;79(3):570–80.

102. Trapasso F, Iuliano R, Boccia A, Stella A, Visconti R, Bruni P, et al. Rat protein tyrosine phosphatase η suppresses the neoplastic phenotype of retrovirally transformed thyroid cells through the stabilization of p27Kip1. Molecular and cellular biology. 2000;20(24):9236–46.

103. Keane MM, Lowrey GA, Ettenberg SA, Dayton MA, Lipkowitz S. The protein tyrosine phosphatase DEP-1 is induced during differentiation and inhibits growth of breast cancer cells. Cancer research. 1996;56(18):4236–43.

104. Balavenkatraman K, Jandt E, Friedrich K, Kautenburger T, Pool-Zobel B, Östman A, et al. DEP-1 protein tyrosine phosphatase inhibits proliferation and migration of colon carcinoma cells and is upregulated by protective nutrients. Oncogene. 2006;25(47):6319–24.

105. Zhang L, Martelli ML, Battaglia C, Trapasso F, Tramontano D, Viglietto G, et al. Thyroid cell transformation inhibits the expression of a novel rat protein tyrosine phosphatase. Experimental cell research. 1997;235(1):62–70.

106. Trapasso F, Yendamuri S, Dumon KR, Iuliano R, Cesari R, Feig B, et al. Restoration of receptor-type protein tyrosine phosphatase η function inhibits human pancreatic carcinoma cell growth in vitro and in vivo. Carcinogenesis. 2004;25(11):2107–14.

107. Kovalenko M, Denner K, Sandström J, Persson C, Groß S, Jandt E, et al. Site-selective dephosphorylation of the platelet-derived growth factor β-receptor by the receptor-like protein-tyrosine phosphatase DEP-1. Journal of Biological Chemistry. 2000;275(21):16219–26.

108. Palka HL, Park M, Tonks NK. Hepatocyte growth factor receptor tyrosine kinase met is a substrate of the receptor protein-tyrosine phosphatase DEP-1. Journal of Biological Chemistry. 2003;278(8):5728–35.

109. Lampugnani MG, Zanetti A, Corada M, Takahashi T, Balconi G, Breviario F, et al. Contact inhibition of VEGF-induced proliferation requires vascular endothelial cadherin, β-catenin, and the phosphatase DEP-1/CD148. The Journal of cell biology. 2003;161(4):793–804.

110. Ding L, Ellis MJ, Li S, Larson DE, Chen K, Wallis JW, et al. Genome remodelling in a basal-like breast cancer metastasis and xenograft. Nature. 2010;464(7291):999–1005.

111. Spring K, Fournier P, Lapointe L, Chabot C, Roussy J, Pommey S, et al. The protein tyrosine phosphatase DEP-1/PTPRJ promotes breast cancer cell invasion and metastasis. Oncogene. 2015.

112. Chabot C, Spring K, Gratton J-P, Elchebly M, Royal I. New role for the protein tyrosine phosphatase DEP-1 in Akt activation and endothelial cell survival. Molecular and cellular biology. 2009;29(1):241–53.

113. Sacco F, Tinti M, Palma A, Ferrari E, Nardozza AP, van Huijsduijnen RH, et al. Tumor suppressor density-enhanced phosphatase-1 (DEP-1) inhibits the RAS pathway by direct dephosphorylation of ERK1/2 kinases. Journal of Biological Chemistry. 2009;284(33):22048–58.

114. Arora D, Stopp S, Böhmer S-A, Schons J, Godfrey R, Masson K, et al. Protein-tyrosine phosphatase DEP-1 controls receptor tyrosine kinase FLT3 signaling. Journal of Biological Chemistry. 2011;286(13):10918–29.

115. Tarcic G, Boguslavsky SK, Wakim J, Kiuchi T, Liu A, Reinitz F, et al. An unbiased screen identifies DEP-1 tumor suppressor as a phosphatase controlling EGFR endocytosis. Current Biology. 2009;19(21):1788–98.

116. Cerami E, Gao J, Dogrusoz U, Gross BE, Sumer SO, Aksoy BA, et al. The cBio cancer genomics portal: an open platform for exploring multidimensional cancer genomics data. Cancer discovery. 2012;2(5):401–4.

117. Gao J, Aksoy BA, Dogrusoz U, Dresdner G, Gross B, Sumer SO, et al. Integrative analysis of complex cancer genomics and clinical profiles using the cBioPortal. Science Signaling. 2013;6(269):1.

118. Veeriah S, Brennan C, Meng S, Singh B, Fagin JA, Solit DB, et al. The tyrosine phosphatase PTPRD is a tumor suppressor that is frequently inactivated and mutated in glioblastoma and other human cancers. Proceedings of the National Academy of Sciences. 2009;106(23):9435–40.

119. Solomon DA, Kim J-S, Cronin JC, Sibenaller Z, Ryken T, Rosenberg SA, et al. Mutational inactivation of PTPRD in glioblastoma multiforme and malignant melanoma. Cancer research. 2008;68(24):10300–6.

120. Walia V, Prickett TD, Kim JS, Gartner JJ, Lin JC, Zhou M, et al. Mutational and Functional Analysis of the Tumor-Suppressor PTPRD in Human Melanoma. Human mutation. 2014;35(11):1301–10.

121. Nair P, DePreter K, Vandesompele J, Speleman F, Stallings RL. Aberrant splicing of the PTPRD gene mimics microdeletions identified at this locus in neuroblastomas. Genes, Chromosomes and Cancer. 2008;47(3):197–202.

122. Stark MS, Woods SL, Gartside MG, Bonazzi VF, Dutton-Regester K, Aoude LG, et al. Frequent somatic mutations in MAP3K5 and MAP3K9 in metastatic melanoma identified by exome sequencing. Nature genetics. 2012;44(2):165–9.

123. Solimini NL, Xu Q, Mermel CH, Liang AC, Schlabach MR, Luo J, et al. Recurrent hemizygous deletions in cancers may optimize proliferative potential. Science. 2012;337(6090):104–9.

124. Goldschmidt MH. Histological classification of epithelial and melanocytic tumors of the skin of domestic animals: Armed Forces Institute of Pathology: American Registry of Pathology: World Health Organization Collaborating Center for Comparative Oncology; 1998.

125. Liang WS, Aldrich J, Tembe W, Kurdoglu A, Cherni I, Phillips L, et al. Long insert whole genome sequencing for copy number variant and translocation detection. Nucleic Acids Res. 2014 Jan;42(2):e8.

126. Li H, Durbin R. Fast and accurate short read alignment with Burrows-Wheeler transform. Bioinformatics. 2009;25(14):1754-60. doi:10.093/bioinformatics/btp324. Epub 2009 May 18.

127. McKenna A, Hanna M, Banks E, Sivachenko A, Cibulskis K, Kernytsky A, et al. The Genome Analysis Toolkit: a MapReduce framework for analyzing next-generation DNA sequencing data. Genome Res. 2010;20(9):1297–303.

128. Van der Auwera GA, Carneiro MO, Hartl C, Poplin R, del Angel G, Levy-Moonshine A, et al. From FastQ data to high-confidence variant calls: the genome analysis toolkit best practices pipeline. Current protocols in bioinformatics. 2013:11.0. 1-.0. 33.

129. Saunders CT, Wong WS, Swamy S, Becq J, Murray LJ, Cheetham RK. Strelka: accurate somatic small-variant calling from sequenced tumor-normal sample pairs. Bioinformatics. 2012;28(14):1811-7. doi:10.093/bioinformatics/bts271. Epub 2012 May 10.

130. Christoforides A, Carpten JD, Weiss GJ, Demeure MJ, Von Hoff DD, Craig DW.Identification of somatic mutations in cancer through Bayesian-based analysis of sequenced genome pairs. BMC Genomics. 2013;14:302.(doi):10.1186/471-2164-14-302.

131. Cibulskis K, Lawrence MS, Carter SL, Sivachenko A, Jaffe D, Sougnez C, et al. Sensitive detection of somatic point mutations in impure and heterogeneous cancer samples. Nat Biotechnol. 2013;31(3):213-9. doi:10.1038/nbt.2514. Epub 013 Feb 10.

132. Cingolani P, Platts A, Wang LL, Coon M, Nguyen T, Wang L, et al. A program for annotating and predicting the effects of single nucleotide polymorphisms, SnpEff: SNPs in the genome of Drosophila melanogaster strain w1118; iso-2; iso-3. Fly. 2012;6(2):80–92.

133. Olshen AB, Venkatraman E, Lucito R, Wigler M. Circular binary segmentation for the analysis of array-based DNA copy number data. Biostatistics. 2004;5(4):557–72.

134. Trapnell C, Pachter L, Salzberg SL. TopHat: discovering splice junctions with RNA-Seq. Bioinformatics. 2009;25(9):1105–11.

135. Angstadt AY, Thayanithy V, Subramanian S, Modiano JF, Breen M. A genome-wide approach to comparative oncology: high-resolution oligonucleotide aCGH of canine and human osteosarcoma pinpoints shared microaberrations. Cancer Genetics. 2012 2012/11//;205(11):572–87.

136. Thomas R, Borst L, Rotroff D, Motsinger-Reif A, Lindblad-Toh K, Modiano JF, et al. Genomic profiling reveals extensive heterogeneity in somatic DNA copy number aberrations of canine hemangiosarcoma. Chromosome Res. 2014;22(3):305–19.

